# Glucose starvation to rapid death of Nrf1, but not Nrf2, deficient hepatoma cells from its fatal defects in the redox metabolism reprogramming

**DOI:** 10.1101/2019.12.13.875369

**Authors:** Yu-ping Zhu, Ze Zheng, Yuancai Xiang, Yiguo Zhang

**Author notes:** Correspondence should be addressed to Yiguo Zhang, or).

## Abstract

Metabolic reprogramming exists in a variety of cancer cells, with the most relevance to glucose as a source of energy and carbon for survival and proliferation. Of note, Nrf1 was shown to be essential for regulating glycolysis pathway, but it is unknown whether it plays a role in cancer metabolic reprogramming, particularly in response to glucose starvation. Herein, we discover that *Nrf1α*^−*/*−^-derived hepatoma cells are sensitive to rapid death induced by glucose deprivation, such cell death appears to be rescued by *Nrf2* interference, but *Nrf1/2*^*+/+*^ or *Nrf2*^−*/*−^-derived cells are roughly unaffected by glucose starvation. Further evidence revealed that *Nrf1α*^−*/*−^ cell death is resulted from severe oxidative stress arising from abberrant redox metabolism. Strikingly, altered gluconeogenesis pathway was aggravated by glucose starvation of *Nrf1α*^−*/*−^ cells, as also accompanied by weakened pentose phosphate pathway, dysfunction of serine-to-glutathione synthesis, and accumulation of reactive oxygen species (ROS) and damages, such that the intracellular GSH and NADPH were exhausted. These demonstrate that glucose starvation leads to acute death of *Nrf1α*^−*/*−^, rather than *Nrf2*^−*/*−^, cells resulting from its fatal defects in the redox metabolism reprogramming. This is owing to distinct requirements of Nrf1 and Nrf2 for regulating key genes involved in the redox metabolic reprogramming. Altogether, this work substantiates the preventive and therapeutic strategies against *Nrf1α*-deficient cancer by limiting its glucose and energy demands.

## Introduction

Metabolic reprogramming is involved in deregulating anabolism and catabolism of glucose, fatty acids and amino acids, which is widely existing in a vast variety of cancer cells [1], so as to facilitate these uncontrolled cell growth and proliferation. Usually, cancer cells increase its uptake of nutrients, mainly including glucose and glutamine. Of note, the ensuing metabolism of glucose, as a major nutrient to fuel cell growth and proliferation, comprises glycolysis pathway, gluconeogenesis pathway, pentose phosphate pathway (PPP), and serine synthesis pathway (SSP), all of which occur in the cytoplasm, besides the tricarboxylic acid cycle (TCA cycle) occurring in the mitochondria [2]. Among these, glycolysis is a central pathway of glucose metabolism but also can be branched towards many anabolic pathways *via* its metabolic intermediary [3]. In cancer cells, decreases in both their oxidative phosphorylation and aerobic glycolysis, are accompanied by increases in the another glycolytic flux, which is independent of oxygen concentration to support the enhanced anabolic demands (of e.g., nucleotides, amino acids and lipids) by providing glycolytic intermediates as raw material [4, 5]. Thereby, such metabolic changes constitute one of typical hallmarks of tumor cells [1, 6].

Clearly, cell life and death decisions are influenced by its cellular metabolism [7], particularly metabolism of cancer cells, which is the most relevant to glucose as a source of energy and carbon. A recent study has uncovered the lower glycolytic rates leading to enhanced cell death by apoptosis [8]. By contrast, the another enforced glycolysis can also effectively inhibit apoptosis [9, 10]. As for the more nutrient uptake than that of normal cells, cancer cells frequently undergo certain metabolic stress due to the shortages in supply of oxygen, nutrients and growth factors. As such, the rapidly proliferating cancer cells were also unable to stop their anabolic and energy requirements, which eventually leads to cell death [11]. Thereby, such a nutrient limitation has been proposed as an effective approach to inhibit the proliferation of cancer cells. For this end, glucose starvation is also considered as a major form of metabolic stress in cancer cells [12]. However, whether (and how) these cell life and death fates are determined in response to metabolic stress induced by glucose starvation remains elusive.

Glucose metabolism is also regulated by the proto-oncogene c-Myc, that was involved in glycolysis by regulating the glycolytic enzymes [13] and also promoted serine biosynthesis upon nutrient deprivation in cancer cells [14]. The another key oncogene HIF-1 was also identified to act as a central regulator of glucose metabolism [15, 16]. Besides, the tumor suppressor p53 can also play a key negative regulatory role in glycolysis by reducing the glucose uptake [17]. Herein, we determine whether (and how) two antioxidant transcription factors Nrf1 (also called Nfe2l1, as a tumor repressor) and Nrf2 (as a tumor promoter) are required for glycolysis and other glucose metabolic pathways, and also involved in the redox metabolic reprogramming induced by glucose deprivation.

Among the cap’n’collar (CNC) basic-region leucine zipper (bZIP) family, Nrf1 and Nrf2 were identified to be two indispensable transcription factors for maintaining redox homoeostasis by binding to antioxidant response elements (AREs) of their downstream target gene promoters [18]. However, recently ever-mounting evidence revealed that the water-soluble Nrf2 activation promotes cancer progression and metastasis [19-21]. Notably, Nrf2 also has a direct or another indirect role in all the hallmarks of cancer, such as mediating metabolic reprogramming [22] and altering redox homeostasis [23]. By contrast, the membrane-bound Nrf1 is subjected to alternative translation and proteolytic processing of this CNC-bZIP protein to yield multiple distinct isoforms of between 140-kDa and 25-kDa; they included TCF11/Nrf1α (120∼140kDa), Nrf1β (∼65kDa), Nrf1γ (∼36kDa), and Nrf1δ (∼25kDa). Among them, Nrf1α was identified to exist as a major isoform in HepaG2 cells as described previously [24]. Specific knockout of Nrf1α (as a dominant tumor repressor) leads to obvious malignant proliferation and tumor metastasis of *Nrf1α*^−*/*−^-derived hepatoma in xenograft model mice [25]. Besides, Nrf1 was also considered to be involved in hepatic lipid metabolism by directly regulating *Lipin1* and *PGC-1* genes [26]. Moreover, Nrf1 was also found to contribute to negative regulation of the Cystine/Glutamate transporter and lipid-metabolizing enzymes [27]. Interestingly, Nrf1 was also positively involved in glycolysis pathway by regulating the *Slc2a2, Gck* [28] and *HK1* genes [29]. However, it is not clear about a role of Nrf1 in mediating the cancer cellular response to metabolic stress, especially stimulated by glucose deprivation.

In this study, we observed a surprising change in the growth of *Nrf1α*^−*/*−^ cells starved in a non-glucose medium. It was found that *Nrf1α*^−*/*−^ cells were more sensitive to glucose deprivation, leading to acute death within 12 h of glucose deprivation, while both cases of *Nrf1/2*^*+/+*^ and *Nrf2*^−*/*−^ were unaffected. As such, the glucose starvation-induced death of *Nrf1α*^−*/*−^ cells was also rescued by *Nrf2* interference. Further examinations revealed that a large amount of reactive oxygen species (ROS) was accumulated by glucose deprivation in *Nrf1α*^−*/*−^ cells, leading to severe oxidative stress. Such a redox imbalance was also attributable to the fact that the intracellular reducing agents (i.e., GSH and NADPH) were exhausted during glucose deprivation. This was due to fatal defects of *Nrf1α*^−*/*−^ cells, resulting in aberrant expression of some key genes (e.g., *CAT, GPX1, GSR, PCK1/2, G6PD, PHGDH* and *ATF4*) that are responsible for the redox metabolism reprogramming of *Nrf1α*^−*/*−^, but not *Nrf2*^−*/*−^ cells, albeit these genes were differentially regulated by Nrf1 and/or Nrf2. Collectively, these demonstrate that Nrf1 and Nrf2 play distinct and even opposing roles in mediating cancer cellular responses to the metabolic stress induced by glucose starvation. Notably, Nrf1 acts as a pivotal player to determine the steady-state level of distinct intracellular redox homeostasis.

## Results

### *Nrf1α*^−*/*−^ cells are susceptible to the cellular death from glucose deprivation

In view of the fact that Nrf1 and Nrf2 have certain overlapping and complementary functions, we here attempted to explore whether and/or how both CNC-bZIP factors exert distinct or opposing roles in mediating the cancer cellular response to metabolic stress. For this, *Nrf1α*^−*/*−^- and *Nrf2*^−*/*−^-deficient HepG2 cell lines (both had been established by Qiu et al [30]), along with their parent wild-type (*Nrf1/2*^*+/+*^) HepG2 cells, were subjected to glucose-free starvation for distinct lengths of time. Subsequently, changes in these cell morphology after glucose deprivation were observed by microscopy. Within 6 h of glucose starvation, no obvious abnormalities of all three cell lines were shown (Figure S1A). In fact, they were growing well, with no changes in the small number of dead cells (Figure S1B). However, when the time of glucose starvation was extended to 12 h, *Nrf1α*^−*/*−^ cells displayed apparent morphological characteristics of cell death (resembling the necrosis and/or necroptosis, Figure 1A), of which such dead cells were stained by trypan blue to 49.26% (Figure 1B). By sharp contrast, both *Nrf1/2*^*+/+*^ and *Nrf2*^−*/*−^ cell lines were largely unaffected by 12 h of glucose starvation (Figure 1, A & B). These cell death or survival was further corroborated by flow cytometry analysis of dual staining cells with fluorescent Annexin V and propidium iodide (PI), showing that *Nrf1α*^−*/*−^ cells were more sensitive (than other two examined cell lines) to cell death induced by glucose deprivation for 12 h (Figure 1C).

**Figure 1.**
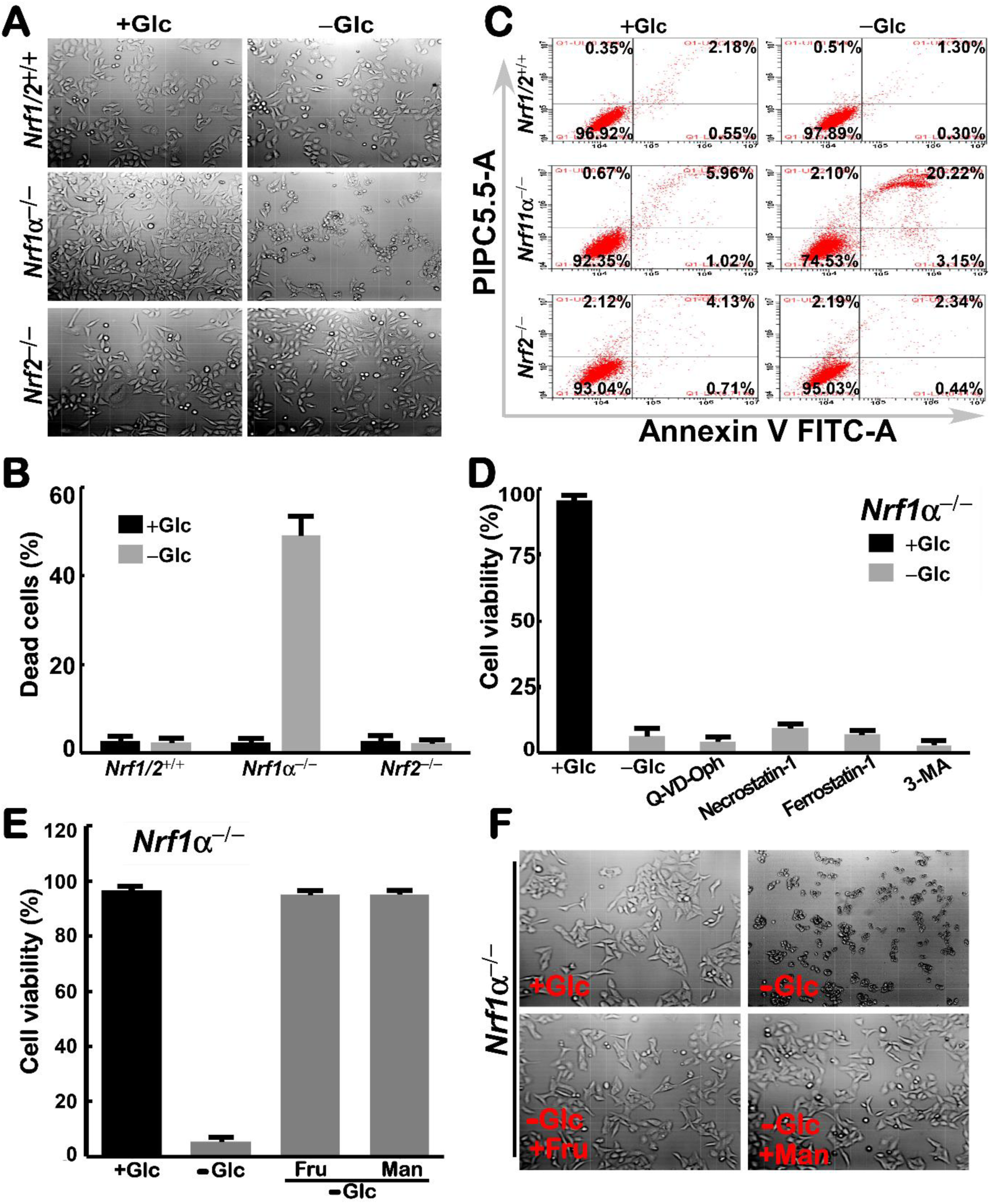
The response of *Nrf1α*^*−/−*^ cells to glucose starvation. (A) Morphological changes of *Nrf1/2*^*+/+*^, *Nrf1α*^*−/−*^ and *Nrf2*^*−/−*^ cells, that had been subjected to glucose deprivation for 12 h, were observed by microscopy (with an original magnification of 200×). (B) The percentage of their dead cells was calculated after being stained by trypan blue. (C) The apoptosis of glucose-starved cells was analyzed by flow cytometry, after being incubated with Annexin V-FITC and PI. (D) *Nrf1α*^*−/−*^ cell viability was determined by incubation for 24 h with q-VD-OPH (10 μM), Necrostatin-1 (100 μM), Ferrostatin-1 (2 μM) or 3-methyladenine (2 mM) in the glucose-free media, each of which was resolved in DMSO as a vehicle. (E) *Nrf1α*^*−/−*^ cell survival was recovered from glucose deprivation by being cultured for 24 h in alternative media containing 25 mM of fructose (Fru) or mannose (Man). (F) Morphology of Fru/Man-recovered *Nrf1α*^*−/−*^ cells was visualized by microscopy (with an original magnification of 200×).

As the glucose starvation was further extended to 24 h, almost all *Nrf1α*^−*/*−^ cells were subjected to the cellular death (Figure S1C). Such prolonged glucose starvation-induced death of *Nrf1α*^−*/*−^ cells was incremented to 95.6% of their examined cells, but only a small number (20%) of *Nrf1/2*^*+/+*^ cells were showed to its cellular death (Figure S1D). Meanwhile, *Nrf2*^−*/*−^ cells appeared to be remaining robust resistant to the putative cellular death stimulated by glucose deprivation for 24 h, however (Figure S1, C & D).

Next, several inhibitors of distinct signaling pathways towards cell death were here employed so as to determine which types of *Nrf1α*^−*/*−^ cell death are resulted from glucose deprivation. Unexpectedly, treatment of *Nrf1α*^−*/*−^ cells with Q-VD-OPH (acting as a pan-caspase inhibitor to block the apoptosis pathway), Necrostatin-1 (as a necroptosis inhibitor), Ferrostatin-1 (as a ferroptosis inhibitor), as well as 3-methyladenine (3-MA, as an autophagy inhibitor), demonstrated that they had not exerted any cytoprotective effects against the cell death caused by glucose deprivation (Figure 1D). Thereby, it is inferable that *Nrf1α*^−*/*−^ cell death from glucose withdrawal may pertain to a major non-apoptotic form of cellular necrosis.

Since glucose deprivation, but not the glycolytic inhibition, leads to death of *Nrf1α*^−*/*−^ cells, we hence investigated whether the cellular death was rescued by other sugar sources, such as fructose or mannose, because both could also serve as a potent alternative to glucose. As shown in Figure 1 (E & F), the results unraveled that fructose and mannose were metabolically utilized in *Nrf1α*^−*/*−^ cells insomuch as to resist against the cell death induced by glucose deprivation. This demonstrates that the lack of sugar source is responsible for determining the death fate of *Nrf1α*^−*/*−^ cells.

### *Nrf1α*^−*/*−^ cell death is driven by glucose deprivation leading to severe endogenous oxidative stress

Clearly, cell life or death fate decisions are selectively determined by the intracellular energy metabolism and redox homeostasis [31]. Thereby, we herein examined whether endogenous oxidative stress of *Nrf1α*^−*/*−^ cells is stimulated by glucose starvation contributing to the cellular responsiveness to death. As anticipated, Figure 2 (A & B) showed that *Nrf1α*^−*/*−^ cell death arising from removal of glucose enabled to be sufficiently rescued after addition of 2-deoxyglucose (2DG, as an analogue of glucose) to the sugar-free culture medium. This is due to the fact that 2DG has a potent ability to inhibit the glycolysis, and thus this treatment of *Nrf1α*^−*/*−^ cells enabled the existing available glucose to enter the PPP route in order to generate certain amounts of NADPH (that acts as a major metabolically reducing agent required for the setting of intracellular basal redox state [32]).

**Figure 2.**
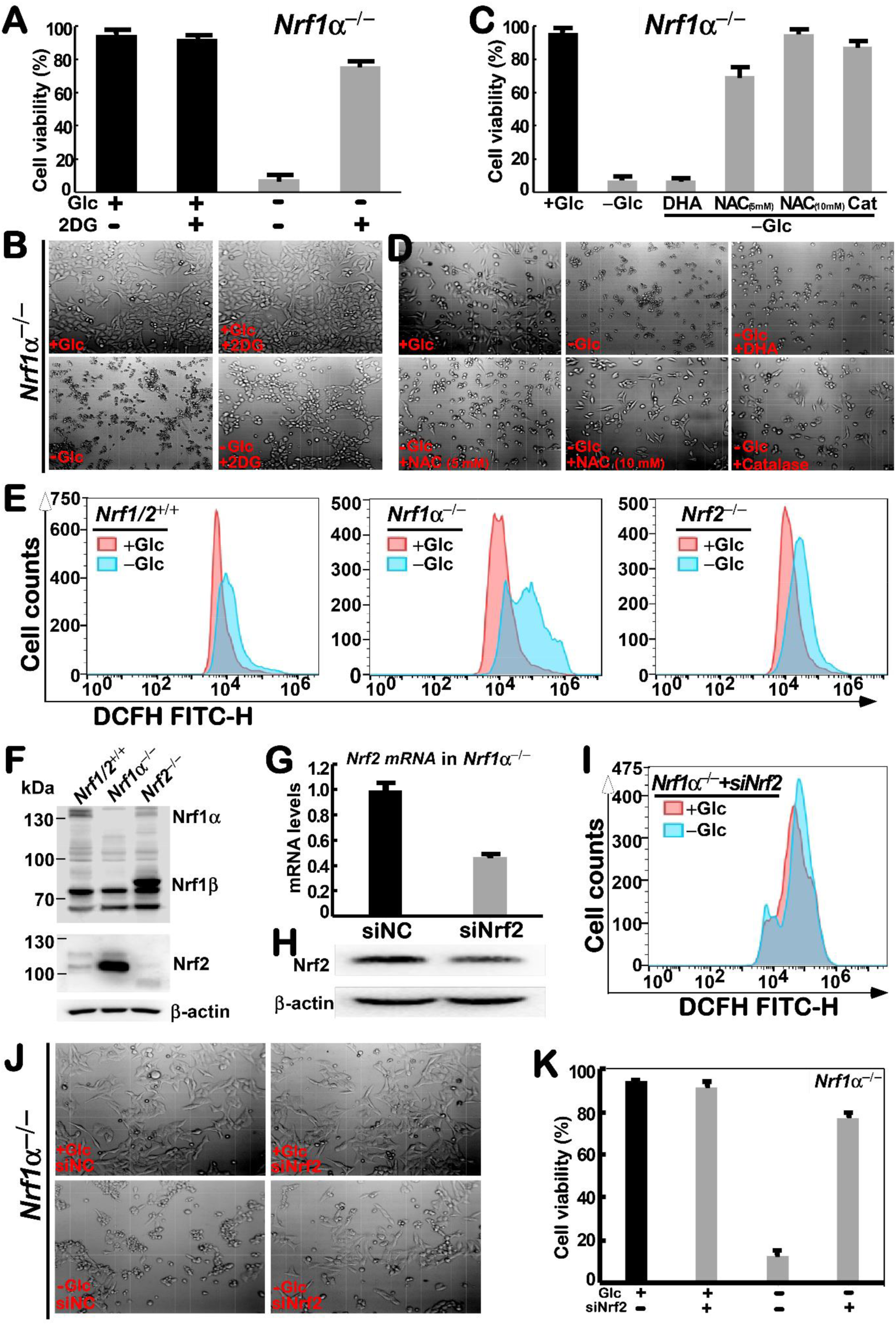
Distinct redox states of starved or rescued *Nrf1α*^*−/−*^ cells with different morphological changes. (A) *Nrf1α*^*−/−*^ cells had been rescued by incubation for 24 h with 2-deoxyglucose (2DG, 10 mM in the glucose-free media), before the cell viability was determined by trypan blue staining. (B) Morphology of 2DG-rescued *Nrf1α*^*−/−*^ cells was visualized by microscopy (with an original magnification of 200×). (C) *Nrf1α*^*−/−*^ cell viability was calculated after they had been cultured for 24 h in glucose-free media containing N-acetyl-cysteine (NAC at 5-10 mM), catalase (CAT at 50 Units/ml) or dehydroascorbic acid (DHA at 100 μM),. (D) Morphological changes of *Nrf1α*^*−/−*^ cells, that had been treated with NAC, CAT or DHA, were observed by microscopy (with an original magnification of 200×). (E) Distinct ROS levels were determined by flow cytometry analysis of *Nrf1/2*^*+/+*^, *Nrf1α*^*−/−*^ and *Nrf2*^*−/−*^ cells that had been starved, or not starved, for 12 h. (F) Abundances of Nrf1 and Nrf2 proteins were determined by Western blotting of *Nrf1/2*^*+/+*^, *Nrf1α*^*−/−*^ and *Nrf2*^*−/−*^ cells. (G, H) The interference of *siNrf2* in *Nrf1α*^*−/−*^ cells was identified by RT-qPCR (*G*) and Western blotting (*H*). (I) Changes in ROS levels of *Nrf1α*^*−/−*^*+siNrf2* cells were analyzed by flow cytometry after 12 h of glucose deprivation. (J) The effects of *siNrf2* on glucose-starved cell morphology were observed by microscopy (with an original magnification of 200×). (K) *Nrf1α*^*−/−*^*+siNrf2* cell viability was evaluated after they had been subjected to glucose starvation for 24 h.

Further examinations revealed that glucose starvation-induced death of *Nrf1α*^−*/*−^ cells was significantly mitigated or completely rescued by treatment with N-acetyl-L-cysteine (NAC, an antioxidant agent to increase glutathione synthesis) or catalase (CAT, an enzyme that catalyzes the breakdown of hydrogen peroxide) (Figure 2, C & D). By contrast, *Nrf1α*^−*/*−^ cell death triggered by glucose starvation was almost unaffected by treatment with dehydroascorbic acid (DHA, as a stable oxidative product of L-ascorbic acid). These suggest that severe oxidative stress may contribute to rapid death of *Nrf1α*^−*/*−^ cells from glucose deprivation. This notion is further evidenced by flow cytometry analysis of distinct cellular oxidative states (Figure 2E). The results unraveled that glucose deprivation caused an obvious accumulation of ROS in *Nrf1α*^−*/*−^ cells, with the oxidative fluorescent images becoming widened and right-shifted, when compared with other two cases of *Nrf1/2*^*+/+*^ and *Nrf2*^−*/*−^ cell lines (only with modestly increased ROS levels) (Figure 2E). Collectively, these imply there exists a fatal defect of *Nrf1α*^−*/*−^, rather than *Nrf2*^−*/*−^, cells in the antioxidant cytoprotective response against the cellular death attack from glucose withdrawal stress.

### Rapid death of glucose-starved *Nrf1α*^−*/*−^ cells can be rescued by interference with *Nrf2*

Based on the similar structure and function of Nrf1 and Nrf2 [33], it is postulated that loss of Nrf1α is likely compensated by Nrf2. As anticipated, Western blotting revealed that an aberrant high expression of Nrf2 was determined in *Nrf1α*^−*/*−^ cells, when compared with that of wild-type *Nrf1/2*^*+/+*^ cells (Figure 2F). Conversely, considerable lower expression levels of Nrf1α-derived protein isoforms were maintained in *Nrf2*^−*/*−^ cells, albeit with a compensatory higher expression of the short Nrf1β (Figure 2F).

Since such hyper-expression of Nrf2 in *Nrf1α*^−*/*−^ cells serves as a complement to *Nrf1α* knockout, it is thus inferable that Nrf2 may also contribute to mediating a putative response of *Nrf1α*^−*/*−^ cells to death attack by glucose starvation. Thereby, we here tried to interfere with the *Nrf2* expression by siRNA transfected into glucose-starved *Nrf1α*^−*/*−^ cells. As shown in Figure 2 (G & H), both mRNA and protein expression levels of Nrf2 were significantly knocked down by its specific siRNA (i.e., *siNrf2*) in *Nrf1α*^−*/*−^ cells. More interestingly, glucose starvation-induced death of *Nrf1α*^−*/*−^ cells was strikingly ameliorated by interference of *siNrf2* (Figure 2, J & K). This is also substantiated by further evidence obtained from flow cytometry analysis of the cellular death (Figure S1E). Such effectively *siNrf2*-alleviated death of *Nrf1α*^−*/*−^ cells was also accompanied by a significant reduction in the intracellular ROS accumulation by glucose deprivation (Figure 2I). Taken together, these observations demonstrate that hyper-active Nrf2 can also make a major contribution to the accumulation of ROS products in *Nrf1α*^−*/*−^ cells, leading to the cellular death driven by glucose starvation.

### The failure of redox defense systems in *Nrf1α*^−*/*−^ cells contributes to its lethality of glucose starvation

The aforementioned evidence (as shown in Figure 2, C to E) demonstrated that glucose starvation-induced death of *Nrf1α*^−*/*−^ cells resulting from severe oxidative stress was effectively prevented by NAC and CAT. This suggests that the intracellular redox state is rebalanced by either NAC or CAT, because NAC facilitates conversion of oxidized glutathione (GSSG) to reduced glutathione (GSH) by glutathione(-disulfide) reductase (GR or GSR), while CAT can catalyze hydrogen peroxide (H_2_O_2_) breakdown to water and oxygen, such that the cytotoxic effects of ROS on *Nrf1α*^−*/*−^ cells are inhibited.

Herein, we further examined changes in basal and glucose starvation-inducible expression of CAT and GSR, as well as other redox cycling and relevant enzymes, including GPX1, PRX1,TRX1, TRX2, NOX4, SOD1 and SOD2 (Figure 3A), in *Nrf1α*^−*/*−^ cells cultured in complete or glucose-free media. As shown in Figure 3, B & C, RT-qPCR analysis of *Nrf1α*^−*/*−^ cells demonstrated significant increases in basal mRNA expression levels of *CAT* and *GPX1* (glutathione peroxidase 1, which catalyzes reduction of H_2_O_2_ and organic hydroperoxides by GSH, so as to protect cells against oxidative damage**s**), when compared to those of *Nrf1/2*^*+/+*^ cells. After glucose withdrawal from the culture of *Nrf1α*^−*/*−^ cells, such basal expression of both *CAT* and *GPX1* was abruptly inhibited close to their levels measured from *Nrf1/2*^*+/+*^ cells (Figure 3, B & C). Similar marked changes in CAT and GPX1 expression were, however, not observed in *Nrf2*^−*/*−^ cells.

**Figure 3.**
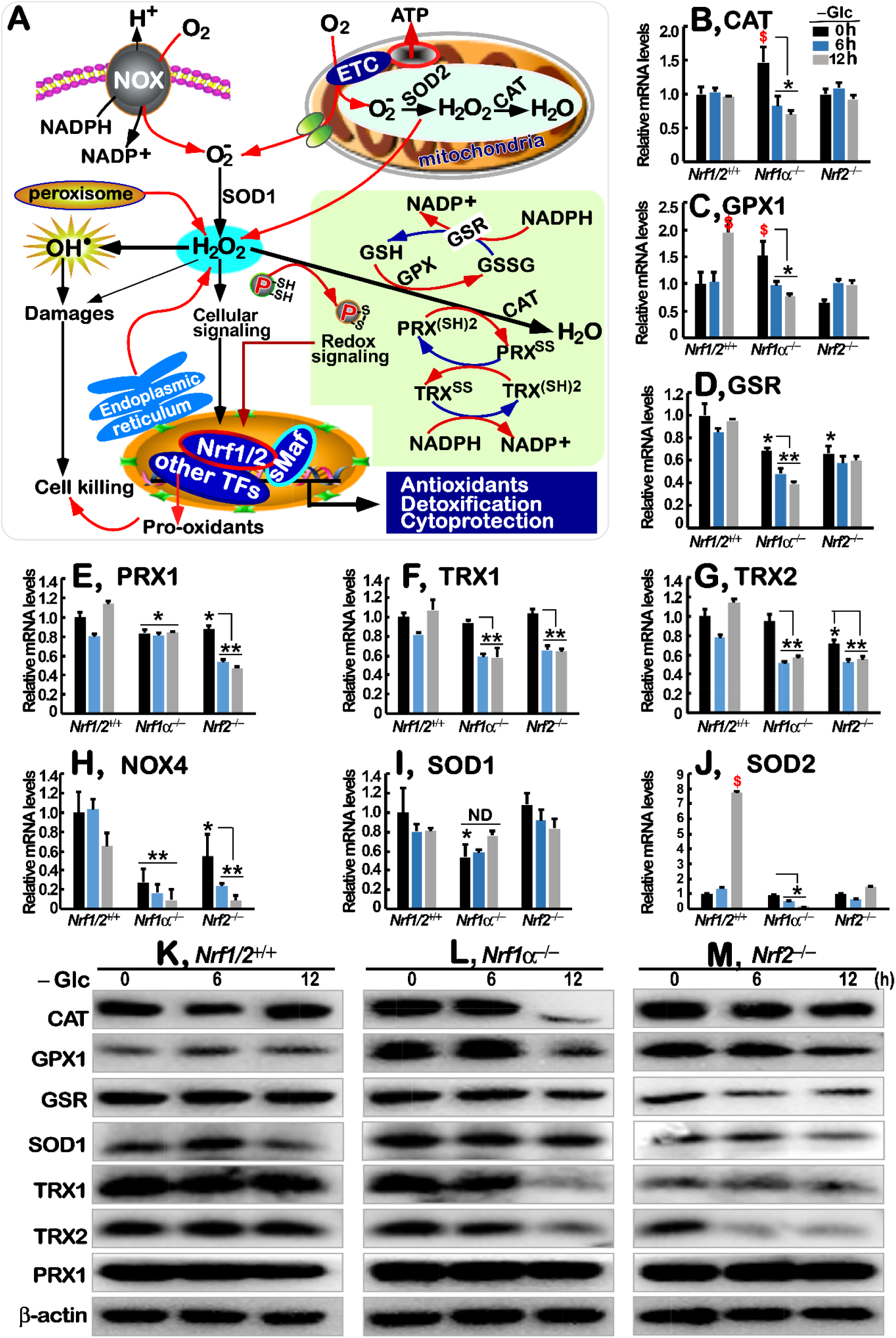
Dysfunction of redox signaling controls and defense systems in glucose-starvated *Nrf1α*^*−/−*^ cells. (A) Schematic representation of intracellular ROS products, along with redox signaling controls and antioxidant defense systems. In this response, Nrf1 and Nrf2 can be induced to translocate the nucleus, in which its functional heterodimer with sMaf or other bZIP proteins is formed in order to transcriptionally regulate distinct subsets of ARE-driven genes, which are responsible for antioxidant, detoxification, and cytoprotection against a variety of cellular stress. (B-J) Alterations in mRNA expression levels of distinct genes, such as (*B*) *GSR* (glutathione disulfide reductase), (*C*) *CAT* (catalase), (*D*) *GPX1* (glutathione peroxidase 1), (*E*) *PRX1* (peroxidase 1), (*F*) *TRX1* (thioredoxin 1), (*G*) *TRX2* (thioredoxin 2), (*H*) *NOX4* (NADPH oxidase 4), (*I*) *SOD1* (superoxide dismutase 1) and (*J*) *SOD2* (superoxide dismutase 2), in *Nrf1/2*^*+/+*^, *Nrf1α*^*−/−*^ and *Nrf2*^*−/−*^ cells, that had been or had not been starved in glucose-free media for 0-12 h, were determined by RT-qPCR analysis. Then, significant increases ($, *p*<0.05) and significant decreases (\**p*<0.05;\*\**p*<0.01) were also calculated, relative to the indicated controls. (K-M) Changes in abundances of the following proteins CAT, GPX1, GSR, SOD1, TRX1, TRX2 and PRX1 were visualized by Western blotting of *Nrf1/2*^*+/+*^ (*K*), *Nrf1α*^*−/−*^ (*L*) and *Nrf2*^*−/−*^ (*M*) cells that had been or had not been glucose-starved for 0-12 h.

By contrast, basal *GSR* expression was down-regulated in *Nrf1α*^−*/*−^ cells, and glucose deprivation also caused it to be further repressed to a considerable lower level, when compared to the wild-type *Nrf1/2*^*+/+*^ cells (Figure 3D). However, similar down-regulation of *GSR* in *Nrf2*^−*/*−^ cells was almost unaffected by glucose starvation. Conversely, expression of *PRX1* (peroxiredoxin-1, also called thioredoxin peroxidase 1) was significantly diminished by glucose deprivation in *Nrf2*^−*/*−^, rather than *Nrf1α*^−*/*−^, cells, albeit its basal expression was similarly down-regulated in these two deficient cell lines (Figure 3E). Further examinations of *Nrf1α*^−*/*−^ and *Nrf2*^−*/*−^ cells revealed that glucose starvation caused remarkable decreases in expression of TRX1 and TRX1 (*i.e.*, both thioredoxin proteins involved in many reversible redox reactions) (Figure 3, F & G). In addition, it is intriguing that deficiency of either *Nrf1α*^−*/*−^ or *Nrf2*^−*/*−^ also led to various extents of decreased expression of *NOX4* (NADPH oxidase 4, which acts as an oxygen sensor and also catalyzes the reduction of molecular oxygen to various ROS), SOD1 (superoxide dismutase 1) and SOD2 (Figure 3, H to I). Notably, only *Nrf1/2*^*+/+*^ cells, but neither *Nrf1α*^−*/*−^ nor *Nrf2*^−*/*−^ cells, showed significant increases in glucose starvation-stimulated expression of *GPX1* and *SOD2* (Figure 3, C & J), but not other examined gene transcripts.

Further Western blotting of *Nrf1α*^−*/*−^ cells unraveled that expression of CAT, GPX1, GSR, TRX1, TRX2, but not PRX1 or SOD1, were substantially decreased by glucose deprivation for 12 h (Figure 3L). By contrast, the abundances of these examined proteins except PRX1 were only marginally decreased to lesser extents by glucose starvation of *Nrf2*^−*/*−^ cells (Figure 3M); such minor effects cannot be excluded to be attributable to a modest decrease of Nrf1α in Nrf2-deficient cells (Figure 2F). In addition, it should be noted that not any increases of the redox-relevant proteins were stimulated by glucose starvation for 6 h to 12 h, even in wild-type *Nrf1/2*^*+/+*^ cells (Figure 3K). Together, it is postulated that *Nrf1α*^−*/*−^, but not *Nrf2*^−*/*−^, cells are much likely to have certain fatal defects in setting the intracellular redox homeostasis along with antioxidant defense systems, such that the resulting redox imbalance contributes to severe endogenous oxidative stress and concomitant damages to *Nrf1α*^−*/*−^ cell death, particularly after 12-h glucose starvation.

### Altered glucose metabolism and energy demands of *Nrf1α*^−*/*−^ cells are deteriorated by glucose deprivation

Since aerobic glycolysis provides the main energy for cancer cells (as illustrated in Figure 4A), it is evitable that the intracellular production of ATP as a major energy source could thus be affected by glucose withdrawal from *Nrf1/2*^*+/+*^, *Nrf1α*^−*/*−^ and *Nrf2*^−*/*−^ cell culture in sugar-free media. As shown in Figure 4B, a substantial diminishment in the basal ATP levels of *Nrf2*^−*/*−^ cells was determined, and even glucose deprivation-stimulated ATP products were also maintained to a considerable lower levels, when compared to those corresponding values measured from *Nrf1/2*^*+/+*^ cells. By contrast, basal ATP levels of *Nrf1α*^−*/*−^ cells (with aberrant hyper-active Nrf2) were significantly elevated (Figure 4B), so as to meet the needs of its malignant growth and proliferation [25, 30]). Intriguingly, such higher ATP production by *Nrf1α*^−*/*−^ cells was further promoted by glucose starvation for 6 h, and thereafter abruptly declined by 12-h prolonged starvation to wild-type basal levels of *Nrf1/2*^*+/+*^ cells (Figure 4B). Accordingly, similar alternations in basal and glucose starvation-stimulated expression of *GLUT1* (glucose transporter 1) were also determined in *Nrf1α*^−*/*−^ cells (Figure 4C), to meet its highly metabolizable energy requirements. This is further supported by Western blotting of *Nrf1α*^−*/*−^ cells displaying a high expression pattern of GLUT1, particularly during glucose-free conditions (Figure 4O). Meanwhile, almost unaltered expression of GLUT1 in *Nrf2*^−*/*−^ cells was observed, even upon glucose starvation (Figure 4P), but this transporter abundances in *Nrf1/2*^*+/+*^ cells were modestly decreased after glucose deprivation (Figure 4N).

**Figure 4.**
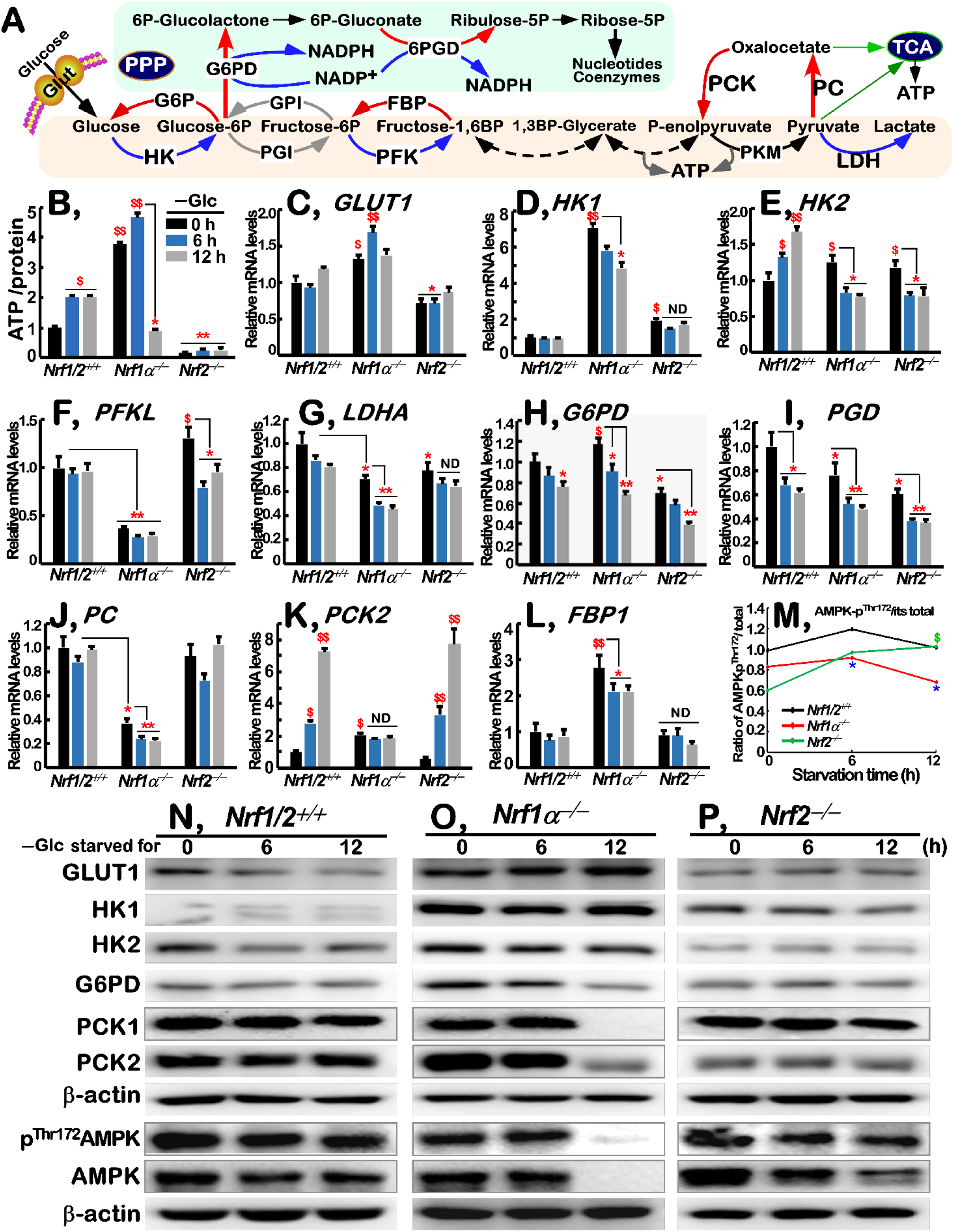
Deterioration of altered glucose metabolism and energy demands by glucose deprivation of *Nrf1α*^*−/−*^ cells. (A) A schematic diagram to give a concise explanation of glycosis, gluconeogenesis and pentose phosphate pathways (PPP). The key rate-limiting enzymes are indicated, apart from TCA (citric acid cycle). In some words, phospho-is represented by a singl *P* letter. (B) Distinct ATP levels of *Nrf1/2*^*+/+*^, *Nrf1α*^*−/−*^ and *Nrf2*^*−/−*^ cells wer determined after glucose deprivation for 0-12 h. (C-L) Altered mRNA expression levels of key metabolic genes: i) (*C*) *GLUT1* (glucose transporter 1), (*D*) *HK1* (hexokinase 1), (*E*) *HK2*, (*F) PFKL* (phosphofructokinase liver type) and (*G*) *LDHA* (lactate dehydrogenase A) involved in the glycolysis pathway; ii) (*H*) *G6PD* (glucose-6-phosphate dehydrogenase) and (I) *PGD* (phosphogluconate dehydrogenase) as rate-limiting enzymes in the PPP; iii) (*J*) *PC* (pyruvate carboxylase), (*K*) *PCK2* (phosphoenolpyruvate carboxykinase 2), and (*L*) *FBP1* (fructose bisphosphatase 1) responsible for the gluconeogenesis pathway, were analyzed by RT-qPCR analysis of *Nrf1/2*^*+/+*^, *Nrf1α*^*−/−*^ and *Nrf2*^*−/−*^ cells that had been starved, or not starved, in the glucose-free media for 0-12 h. Then, significant increases ($, *p*<0.05; $$, *p*<0.01) and significant decreases (\**p*<0.05;\*\**p*<0.01) were also calculated, relative to the indicated controls. (M) Phospho-AMPK^Thr172^/AMPK ratios were calculated by the intensity of their immunoblots in *Nrf1/2*^*+/+*^, *Nrf1α*^*−/−*^ and *Nrf2*^*−/−*^ cells. (P-R) Changes in protein abundances of GLUT1, HK1, HK2, G6PD, PCK1, PCK2, AMPK and its phospho-AMPK^Thr172^ were determined by Western blotting of *Nrf1/2*^*+/+*^ (*P*), *Nrf1α*^*−/−*^ (*Q*) and *Nrf2*^*−/−*^ (*R*) cells, after having been glucose-starved, or not, for 0-12 h.

Besides GLUT1, other key metabolic enzymes (e.g., HK1/2, PFKL and LDHA) required for the glycolysis pathway of cancer cells were also investigated herein. Among them, only hexokinase 2 (HK2) was transcriptionally activated by glucose starvation of *Nrf1/2*^*+/+*^ cells, but not *Nrf1α*^−*/*−^ or *Nrf2*^−*/*−^ cells (Figure 4F). Although basal mRNA expression of *HK2* was obviously up-regulated, but rather its glucose starvation-stimulated expression was significantly suppressed, in *Nrf1α*^−*/*−^ or *Nrf2*^−*/*−^ cells (Figure 4F). Similarly, a substantial increase in basal *HK1* expression occurred in *Nrf1α*^−*/*−^ cells, but its transcriptional expression was strikingly reduced by glucose deprivation (Figure 4E). By contrast, *Nrf2*^−*/*−^ cells also showed a considerable high level of basal *HK1* expression, albeit its mRNA transcription was unaffected by glucose starvation. Further examinations by Western blotting revealed that both HK1 and HK2 protein expression levels were modestly decreased by glucose starvation of *Nrf1α*^−*/*−^ cells, but appeared to be unaffected in glucose-starved *Nrf1/2*^*+/+*^ and *Nrf2*^−*/*−^ cells (Figure 4, N to P). Moreover, apparent decreases in basal *PFKL* (phosphofructokinase, liver type) and *LDHA* (lactate dehydrogenase A) expression levels were observed in *Nrf1α*^−*/*−^ cells (Figure 4, F & G), of which the latter *LDHA* expression was further decreased by glucose starvation, while *PFKL* was only slightly reduced by this stimulation. Contrarily, *Nrf2*^−*/*−^ cells showed a substantial increment in basal *PFKL* expression, but its mRNA transcription expression was markedly suppressed upon glucose deprivation (Figure 4F). By comparison, a modest decrease in basal expression of *LDHA* was also observed in *Nrf2*^−*/*−^ cells, but its expression was roughly unaffected (Figure 4G). These collective data demonstrate distinct roles of Nrf1 and Nrf2 in controlling expression of those key genes responsible for glycolysis.

Since the above observation (Figure 2, A & B) uncovered that glucose-starved *Nrf1α*^−*/*−^ cell death was rescued by 2DG, this glucose analogue could render the metabolic flow to enter the PPP and hence promote NADPH products. As anticipated, RT-qPCR analysis of the rate-limiting enzymes of the PPP showed that mRNA expression levels of *G6PD* (glucose-6-phosphate dehydrogenase) and *PGD* (phosphogluconate dehydrogenase) were significantly decreased after glucose deprivation of *Nrf1/2*^*+/+*^, *Nrf1α*^−*/*−^ and *Nrf2*^−*/*−^ cells (Figure 4, H & I), albeit directionally positive and negative regulation of their basal expression by *Nrf1α*^−*/*−^ or *Nrf2*^−*/*−^ was also determined. Further Western blotting also unraveled that G6PD protein abundances were significantly decreased in glucose-starved *Nrf1α*^−*/*−^, rather than *Nrf1/2*^*+/+*^ and *Nrf2*^−*/*−^, cells (Figure 4, N-P).

From the aforementioned evidence, it is postulated that the gluconeogenesis pathway should be enhanced after glucose deprivation, so that the resulting products were allowed to enter the PPP and other metabolic pathways. Thus, we investigated expression of certain key enzymes involved in the gluconeogenesis. As shown in Figure 4K, a significant increment in transcriptional expression of *PCK2* (phosphoenolpyruvate carboxykinase 2) in *Nrf1/2*^*+/+*^ and *Nrf2*^−*/*−^ cells was stimulated by glucose starvation for 6 h to 12 h. However, no changes in mRNA expression of *PCK2* were detected in glucose-starved *Nrf1α*^−*/*−^ cells, although its basal expression was up-regulated (Figure 4K). This is further supported by Western blotting of PCK1 and PCK2, revealing that both protein abundances were significantly decreased by glucose starvation, especially for 12 h, of *Nrf1α*^−*/*−^ cells, but rather almost unaffected in glucose-starved *Nrf1/2*^*+/+*^ and *Nrf2*^−*/*−^ cells (Figure 4, N-P). Further examinations of both *PC* (pyruvate carboxylase) and *FBP1* (fructose-bisphosphatase 1) unraveled that their mRNA levels were markedly reduced by glucose deprivation in *Nrf1α*^−*/*−^ cells, but largely unaltered in glucose-starved *Nrf1/2*^*+/+*^ or *Nrf2*^−*/*−^ cell lines (Figure 4, J & L). In addition, it should also be noted that *Nrf1α*^−*/*−^ cells manifested down-regulation of basal *PC* expression, along with up-regulation of basal *FBP1* expression.

Next, we examined changes in active phosphorylation of AMPK (AMP-activated protein kinase), since it acts as a key regulator of energy metabolism [34]. As illustrated in Figure 4M, the ratio of phosphorylated AMPK ^Thr172^ to its total protein (as calculated by stoichiometry of their immunoblots as shown in Figure 4, N-P) was significantly increased by glucose withdrawal from *Nrf2*^−*/*−^ cells, but conversely decreased in glucose-starved *Nrf1α*^−*/*−^ cells. The latter notion was also further supported by the fact that almost all of total AMPK and its phospho-AMPK^Thr172^ proteins were evidently abolished by 12-h glucose starvation of *Nrf1α*^−*/*−^ cells (Figure 4O), besides their significant reduction in glucose-starved *Nrf2*^−*/*−^ cells (Figure 4P). Overall, the inactivation of cellular energy switch, along with blockage of its gluconeogenesis and ablation of both its PPP and glycolysis, is inferable as a crucial determinant of *Nrf1α*^−*/*−^ cell death, resulting from glucose deprivation to deteriorate its altered energy metabolic demands.

### Up-regulation of serine-to-glutathione synthesis by glucose deprivation is fatally abolished in *Nrf1α*^−*/*−^ cells, leading to severe endogenous oxidative stress

As illustrated in Figure 5A, *de novo* serine synthesis from the glycolytic metabolite 3-phosphoglycerate (3PG) by SSP, contributes to major carbons for glutathione biosynthesis and also provides precursors for purine and pyrimidine biosynthetic pathway *via* the folate cycle. In the successive biochemical course, 3PG, as a key intermediate of glycolytic pathway, is allowed to flow into the SSP and thus limit ATP production, while oxidation of 3PG converts NAD^+^ to NADH so as to affect intracellular redox state. As a result, serine can also be converted to cysteine and glycine by key enzymes (Figure 4A), for maintaining intracellular glutathione homeostasis. Therefore, we herein investigated putative effects of glucose deprivation on serine-to-glutathione synthesis pathways.

**Figure 5.**
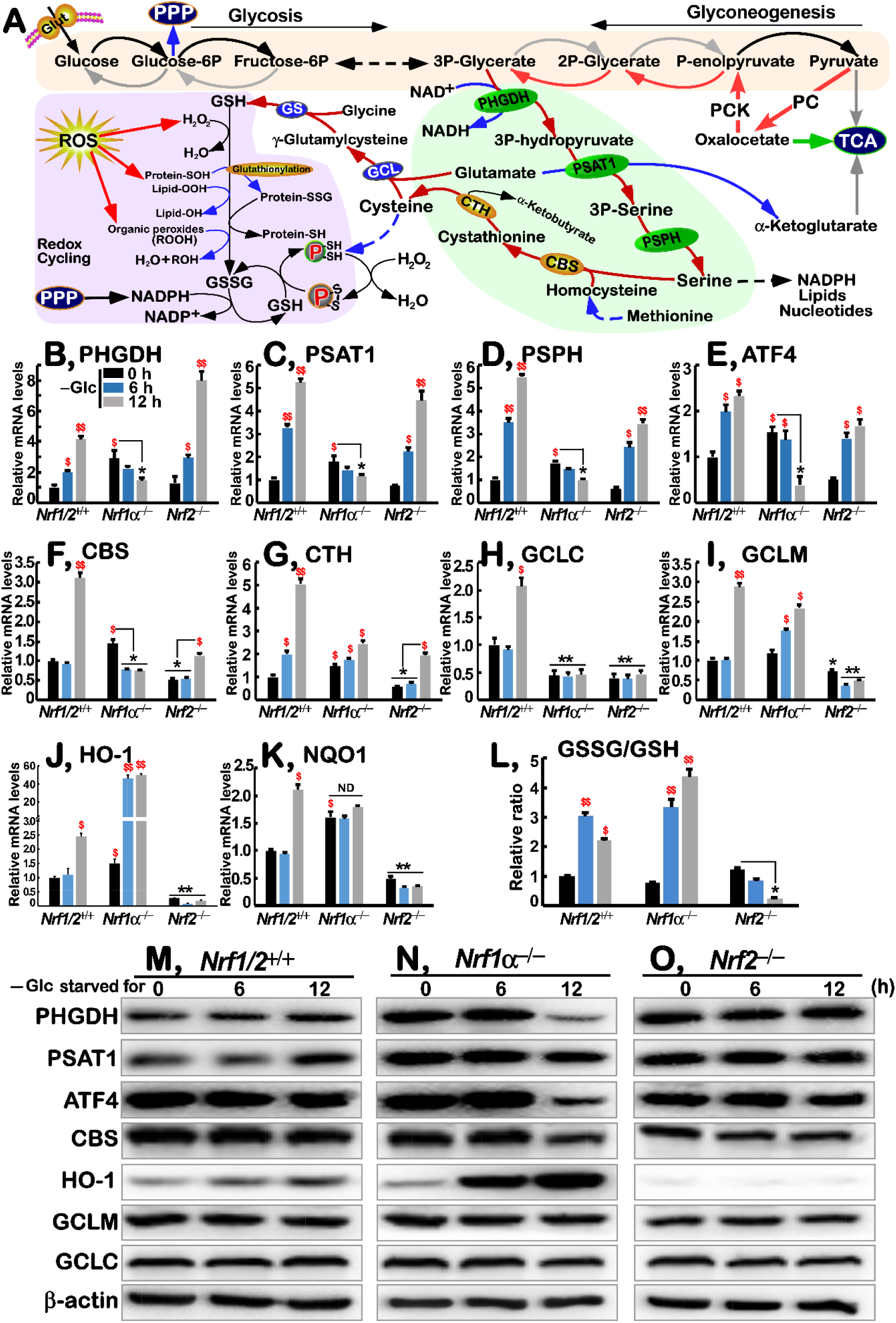
Fatal abolishment of *de novo* serine-to-glutathione biosynthesis by glucose deprivation of *Nrf1α*^*−/−*^ cells. (A) A schematic to give a concise explanation of *de novo* serine synthesis pathway (SSP), along with ensuing transsulfuration to yield cysteine and glutathione (GSH). Both major buffers of GSH/GSSG and NADPH/NADP^+^ are tightly regulated by redox cycling switches and relevant defense systems against ROS and oxidative damages. (B-K) Altered mRNA expression levels of key biosynthetic genes as follows: i) (*B*) *PHGDH* (phosphoglycerate dehydrogenase), (*C*) *PSAT1* (phosphoserine aminotransferase 1), (*D*) *PSPH* (phosphoserine phosphatase) involved in the SSP, along with its regulator (*E*) *ATF4* (activating transcription factor 4); ii) (*F*) *CBS* (cystathionine beta-synthase) and (*G*) *CTH* (cystathionine gamma-lyase) essential for the transsulfuration to yield cysteine; iii) (*H*) *GCLC* (glutamate-cysteine ligase catalytic subunit) and (*I*) *GCLM* (glutamate-cysteine ligase modifier subunit) to catalyze glutathione biosynthesis; iv) as well as antioxidant genes, such as (*J*) *HO-1* (heme oxygenase 1) and (*K*) *NQO1* (NAD(P)H quinone dehydrogenase 1), were determined by RT-qPCR of *Nrf1/2*^*+/+*^, *Nrf1α*^*−/−*^ and *Nrf2*^*−/−*^ cells that had been starved, or not starved, in the glucose-free media for 0-12 h. Then, significant increases ($, *p*<0.05; $$, *p*<0.01) and significant decreases (\**p*<0.05;\*\**p*<0.01) were also calculated, relative to the indicated controls. (L) Effects of glucose deprivation on the intracellular GSSG/GSH ratios in *Nrf1/2*^*+/+*^, *Nrf1α*^*−/−*^ and *Nrf2*^*−/−*^ cells were assessed. (M-O) Changed abundances of PHGDH, PSAT1, ATF4, CBS, HO-1, GCLM and GCLC proteins in *Nrf1/2*^*+/+*^ (*M*), *Nrf1α*^*−/−*^ (*N*) and *Nrf2*^*−/−*^ (*O*) cells were visualized by Western blotting after glucose deprivation for 0-12 h.

As expected, significant increases in mRNA expression of those rate-limiting enzymes, such as *PHGDH* (phosphoglycerate dehydrogenase), *PSAT1* (phosphoserine aminotransferase 1) and *PSPH* (phosphoserine *phosphatase), required in the SSP, as well as their upstream regulatory factor ATF4* (activating transcription factor 4) [35], were triggered by glucose deprivation in *Nrf1/2*^*+/+*^ and *Nrf2*^−*/*−^ cells (Figure 5, B-E). By sharp contrast, transcriptional induction of *PHGDH, PSAT1, PSPH* and *ATF4* by glucose deprivation was completely blocked or suppressed to lesser extents in starved *Nrf1α*^−*/*−^ cells, albeit with an exception of evident increases in their basal mRNA expression up-regulated by loss of *Nrf1α* (Figure 5, B-E). Accordingly, Western blotting showed that glucose starvation of *Nrf1/2*^*+/+*^ cells for 12 h led to modest increases in abundances of PHGDH and PSAT1, rather than ATF4, to greater extents (Figure 5M). However, all three protein expression levels were strikingly decreased by glucose deprivation in *Nrf1α*^−*/*−^ cells, but unaltered in glucose-starved *Nrf2*^−*/*−^ cells (Figure 5, N & O).

Further insights into cysteine metabolism by transsulfuarion enzymes, such as CBS (cystathionine β-synthase) and CTH (cystathionine γ-lyase), revealed that both were transcriptionally up-regulated by glucose starvation in *Nrf1/2*^*+/+*^ cells (Figure 5, F & G). Similarly, *Nrf2*^−*/*−^ cells also manifested modest induction of *CBS* and *CTH* by glucose deprivation, notwithstanding down-regulation of their basal expression by loss of Nrf2 (Figure 5, F & G). Conversely, up-regulation of basal *CBS* and *CTH* expression occurred in *Nrf1α*^−*/*−^ cells (with hyper-active Nrf2). However, glucose starvation triggered oppposing effects on transcriptional expression of *CBS* and *CTH* in *Nrf1α*^−*/*−^ cells; the former *CBS* mRNA levels were evidently suppressed, while the latter *CTH* mRNA expression was marginally induced (Figure 5, F & G). In addition, glucose deprivation also led to an obvious decrease in CBS protein levels in *Nrf1α*^−*/*−^ or *Nrf2*^−*/*−^ cells, when compared to its abundances measured from *Nrf1/2*^*+/+*^ cells (Figure 5, M-O).

Next, effects of glucose deprivation on glutamate-cysteine ligase catalytic and midifier subunits (GCLC and GCLM, both comprise a key rate-limiting enzyme of glutathione biosynthesis) was investigated here. As anticipated, both *GCLC* and *GCLM* mRNA expression was significantly induced by glucose deprivation of *Nrf1/2*^*+/+*^ cells for 12 h (Figure 5H & I). Such induction of *GCLC* expression by glucose deprivation was completely abolished in *Nrf1α*^−*/*−^ or *Nrf2*^−*/*−^ cells, in which a considerable lower basal expression of *GCLC* was maintained by comparison with wild-type levels of *Nrf1/2*^*+/+*^ cells (Figure 5H). By contrast, basal and glucose starvation-stimulated *GCLM* expression levels were up-regulated in *Nrf1α*^−*/*−^ cells, but rather down-regulated in *Nrf2*^−*/*−^ cells (Figure 5I). Howerver, *GCLC* and *GCLM* protein levels were almost unaffected after glucose starvation of the above examined three cell lines (Figure 5, M-O).

Besides *GCLC* and *GCLM*, both *HO-1* (heme oxygenase 1, also called HMOX1) and *NQO1* (NAD(P)H quinone dehydrogenase 1) serves as downstream antioxidant genes of Nrf2 [36]. Here, investigation by RT-qPCR revealed that glucose starvation for 12 h caused significant induction of *HO-1* and *NQO1* expression in *Nrf1/2*^*+/+*^ cells (Figure 5, J & K). By comparison, basal mRNA expression levels of *HO-1* and *NQO1* were up-regulated in *Nrf1α*^−*/*−^ cells, but rather down-regulated in *Nrf2*^−*/*−^ cells. Interestingly, remarkable induction of *HO-1*, rather than *NQO1*, by glucose deprivation was determined in *Nrf1α*^−*/*−^ cells (Figure 5, J & K). Conversely, transcriptional induction of *HO-1* and *NQO1* by glucose deprivation was roughly abolished or even slightly repressed in *Nrf2*^−*/*−^ cells. Furthermore, Western blotting showed that HO-1 protein abundances were significantly induced by glucose starvation of *Nrf1/2*^*+/+*^ and *Nrf1α*^−*/*−^ cells for 6-12 h, but appeared to be completely abolished in *Nrf2*^−*/*−^ cells (Figure 5, M-O). Moreover, further examination unraveled that glucose deprivation caused a significant increase in the GSSG/GSH ratio in *Nrf1/2*^*+/+*^ or *Nrf1α*^−*/*−^ cells, but this ratio was reversively decreased in *Nrf2*^−*/*−^ cells (Figure 5L). Together, these findings demonstate that death of *Nrf1α*^−*/*−^ cells is a consequence of severe endogenous oxidative stress and damages induced by glucose starvation. This is attributable to fatal defects of *Nrf1α*^−*/*−^ cells in the redox metabolic reprogramming, such that the intracellular GSH/GSSG imbalance is further deteriorated by glucose deprivation, even though certain antioxidant response genes are abberrantly activated by hyper-expression of Nrf2 in *Nrf1α-*deficient cells.

### Distinct requirements of Nrf1 and Nrf2 for the redox metabolic reprogramming in response to glucose deprivation

To determine distinct roles of Nrf1 and Nrf2 in mediating cellular redox metabolic responses to glucose deprivation, we examined the expression of Nrf1 and Nrf2 *per se* after glucose starvation of different genotypic cells for 6-12 h. The RT-qPCR showed that transcriptional expression of *Nrf1* and *Nrf2* was significantly induced by glucose deprivation in *Nrf1/2*^*+/+*^ cells (Figure 6, A & B). However, such inducible mRNA expression levels of *Nrf1* and *Nrf2* by glucose starvation, as well as their basal expression, were almost completely abolished in *Nrf1α*^−*/*−^ and *Nrf2*^−*/*−^ cells, respectively (Figure 6, A & B). Interestingly, glucose starvation of *Nrf2*^−*/*−^ cells for 12 h also caused a modest increase in *Nrf1* mRNA expression (Figure 6A), even though evident decreases in its basal mRNA levels (Figure 6A) and Nrf1α-derived proteins (Figure 2F) were determined. Conversely, *Nrf1α*^−*/*−^ cells only manifested a modest induction of *Nrf2* mRNA expression by glucose deprivation, but with no changes in its basal mRNA levels (Figure 6B), albeit its proteins were strikingly accumulated by loss of Nrf1α (Figure 2F). Collectively, these demonstrate bi-directional inter-regulatory roles of between Nrf1 and Nrf2 in distinct contributions to the cellular response triggered by glucose deprivation, in which Nrf1α is a dominant player.

**Figure 6.**
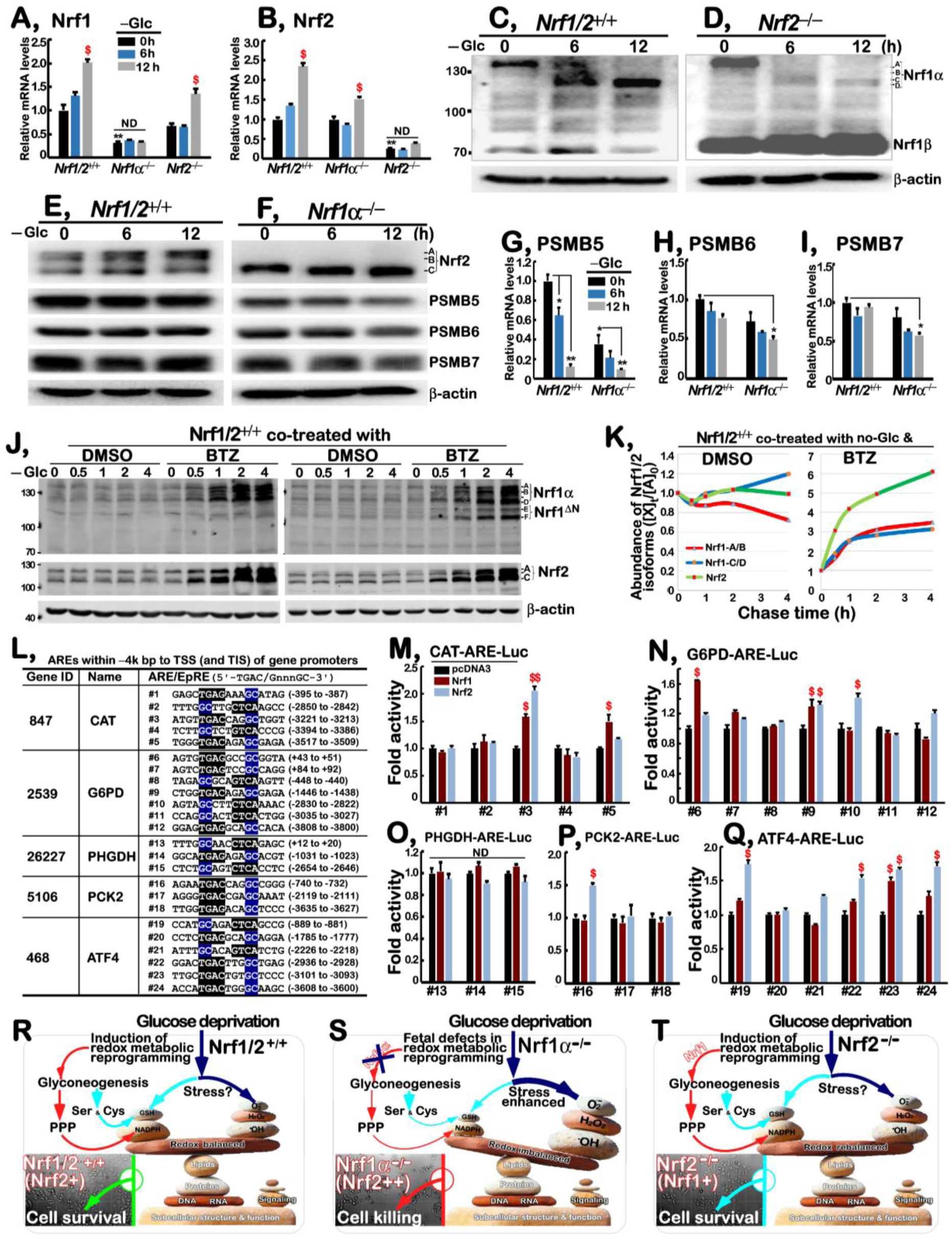
Distinct requirements of Nrf1 and Nrf2 for the cytoprotective response to glucose deprivation. (A,B) Distinct mRNA levels of *Nrf1* (*Nfe2l1*) (*A*) and *Nrf2* (*Nfe2l2*) (*B*) were determined by RT-qPCR analysis of *Nrf1/2*^*+/+*^, *Nrf1α*^*−/−*^ and *Nrf2*^*−/−*^ cells, that had been starved, or not starved, for 0-12 h in the glucose-free media. Then, significant increases ($, *p*<0.05) and significant decreases (\*\**p*<0.01) were calculated, relative to the indicated controls. ND, non-significant difference. (C,D) Changes in Nrf1-derived protein isoforms in *Nrf1/2*^*+/+*^ (*C*) and *Nrf2*^*−/−*^ (*D*) cells were visualized by Western blotting after glucose deprivation for 0-12 h. (E,F) Western blotting of Nrf2, PSMB5, PSMB6 and PSMB7 proteins in *Nrf1/2*^*+/+*^ (*E*) and *Nrf1α*^*−/−*^ cells (*F*) was conducted after 0-12 h of glucose deprivation. Significant decreases (\**p*<0.05; \*\**p*<0.01) were then calculated, relative to the indicated controls. (G-I) Alterations in mRNA levels of *PSMB5* (*G*), *PSMB6* (*H*) and *PSMB7* (*I*) in *Nrf1/2*^*+/+*^ and *Nrf1α*^*−/−*^ cells were determined after glucose starvation for 0-12 h. (J) *Nrf1/2*^*+/+*^ cells were or were co-treated for 0-4 h in the glucose-free media containing 1 µmol/L bortezomib (BTZ, a proteasomal inhibitor) or 0.1% DMSO vehicle, followed by Western blotting with antibodies against Nrf1 or Nrf2. (K) The intensity of the immunoblots representing distinct Nrf1-derived isoforms or Nrf2 proteins, respetively, in the above-treated *Nrf1/2*^*+/+*^ cells (*J*) was quantified by the Quantity One 4.5.2 software, and then shown graphically. (L) 24 of the indicated ARE-adjoining sequences searched from the promoter regions of *CAT, G6PD, PHGDH, PCK2* and *ATF4* were cloned into the pGL3-Promoter vector, and the resulting contrusts served as *ARE-*driven luciferase (*ARE*-*Luc*) reporter genes. (M-Q) *Nrf1/2*^*+/+*^ HepG2 cells were co-transfected with each of the above indicated *ARE-Luc* or non-*ARE-Luc* (as a background control) plasmids, together with an expression construct for Nrf1, Nrf2 or empty pcDNA3.1 vector, then allowed for 24-h recovery before the luciferase activity measured. The results were calculated as a fold change (mean±S.D, n=9) of three independent experiments. Then, significant increases ($, *p*<0.05) were also evaluated, relative to the indicated controls. ND, non-significant difference. (R-T) Three distinct models are proposed to provide a better understanding of molecular basis for survival or death decisions made by glucose-starved *Nrf1/2*^*+/+*^ (*R*), *Nrf1α*^*−/−*^ (*S*) and *Nrf2*^*−/−*^ (*T*) cells. In redox metabolic reprogramming caused by glucose deprivation, the glycosis was diminished or abolished, and thus replaced by increased glyconeogenesis. As a result, many of their intermediates are diverted to enter the PPP and serine-to-glutathione biosynthesis pathways, in order to yield certain amounts of GSH and NADPH. These two reducing agents enable cytoprotective adaptation to oxidative stress induced by glucose deprivation (*R*). However, rapid death of *Nrf1α*^*−/−*^ cells results from its fatal defects in the redox metabolic reprogramming in cellular response to glucose starvation, as accompanied by severe oxidative stress and damage accumulation (*S*). Thereby, Nrf1 is reasonable as a dominant player in the key gene regulation of redox metabolic reprogramming caused by glucose deprivation. As a result, the existence of Nrf1 in *Nrf2*^*−/−*^ cells can still endow their survival with its redox metabolic reprogramming in rebalanced redox state (*T*).

Intriguingly, Nrf1 and Nrf2 exhibited two different but similar trends in their protein levels. Glucose starvation of *Nrf1/2*^*+/+*^ or *Nrf2*^−*/*−^ cells stimulated conversion of Nrf1 glycoprotein-A and then proteolytic processing to yield mature cleaved protein-C/D isoforms before transcriptionally regulating target genes. Thereby, Figure 6 (C & D) showed that glycoprotein-A of Nrf1 was gradually disappeared from 6 h to 12 h in glucose-starved *Nrf1/2*^*+/+*^ or *Nrf2*^−*/*−^ cells, and then was replaced by gradual enhancement of active cleaved Nrf1 protein-C/D. By constrat, Nrf2 proteins in *Nrf1/2*^*+/+*^ or *Nrf1α*^−*/*−^ cells were margnally increased by glucose starvation for 6-12 h (Figure 6, E & F). Such distinct abundances in both Nrf1 and Nrf2 proteins, together with both discrapant mRNA expression levels (Figure 6, A & B), are attributable to their distinct stability and trans-activity during glucose deprivation.

Given that Nrf1, but not Nrf2, exerts an essential biological role in transcriptional expression of proteasomes (PSM), we hence examined potenatial effects of glucose deprivation on Nrf1-target *PSM* genes, to gain a better understanding of disparate contributions of Nrf1 and Nrf2 to death of *Nrf1α*^−*/*−^, but not *Nrf2*^−*/*−^, cells suffered from glucose starvation. As anticipated, both mRNA and protein levels of *PSMB5, PSMB6* and *PSMB7* (encoding the core enzymatic active β5, β1, and β2 subunits, respectively) were significantly abolished or suppressed in glucose-starved *Nrf1α*^−*/*−^ cells, besides down-regulation of their basal expression levels to varying extents (Figure 6, E-I). Such being the case, all three core subunits *PSMB5, PSMB6* and *PSMB7* in *Nrf1/2*^*+/+*^ cell were also not induced by glucose deprivations. Reversively, mRNA expression of *PSMB5* in *Nrf1/2*^*+/+*^ cells was significantly suppressed to less than 20% of its basal level during glucose starvation from 6 h to 12 h, but with no obvious changes in expression of *PSMB6* and *PSMB7* (Figure 6, G-I). Altogether, these demonstrate that negative regulation of proteasomal expression by glucose deprivation is much likely to result in accumulation of oxidative damaged proteins, particularly in glucose-starved *Nrf1α*^−*/*−^ cells.

To clarify distinct contributions of Nrf1 and Nrf2 to mediating cellular responses induced by glucose starvation, we determined conversion of these two CNC-bZIP proteins-derived isoforms and their stability during glucose deprivation As shown in Figure 6J, time-course analysis revealed that the full-length Nrf1α glycoprotein-A was gradually converted into deglycoprotein-B, and ensuing processed protein-C/D in glucose-starved *Nrf1/2*^*+/+*^ cells (of note, similar processing of Nrf1 had been interpreted in details, as elsewhere [37]). Consequently, Nrf1α glycoprotein-A and transient deglycoprotein-B became gradully fainter within 4 h of glucose starvation, while its protein-C/D abundances were conversely enhanced (Figure 6K, *left panel*). All these Nrf1α-derived isoforms were accumulated by co-treatment with the proteasomal inhibitor bortezomib (BTZ) (Figure 6, J & K). Furthermore, stability of Nrf1 precursor protein-A/B and its mature processed protein-C/D in glucose-starved *Nrf1/2*^*+/+*^ cells was estimated by their distinct halflives, which were determined to be 0.24 h (=14.4 min) and 2.53 h (= 151.8 min), respectively, after treatment with cycloheximide (CHX) (Figure S2). Of note, even in the presence of BTZ, glucose deprivation stimulated a rapid processing mechanism of Nrf1α glycoprotein-A and deglycoprotein-B (with a collective halflife of 0.41 h = 24.6 min) to yield certain amonts of its processed protein-C/D (with a more than 4-h halflife, Figures 6J & S2). By contrast, Nrf2 protein levels were almost unaffected by glucose starvation of *Nrf1/2*^*+/+*^ cells, but their abundances were further enhanced by BTZ (Figure 6, J & K). Moreover, Nrf2 potein stabiliy under glucose deprivation conditions was determined by its halflife, that was estimated to be 0.42 h (= 25.2 min) after CHX treatment, but also extended by BTZ to 1.10 h (= 66 min) (Figure S2). Thereby, it is inferable that discrepant stability of Nrf1 and Nrf2 is dedicated to distinct roles of both CNC-bZIP factors in mediating disparate cellular responses to glucose starvation.

Next, to gain insights into direct roles of Nrf1 and Nrf2 in mediating key gene transcriptional responses required for redox metabolic reprogramming, we established 24 of the indicated luciferase reporter genes driven by consensus ARE sequences from the *CAT, G6PD, PHGDH, PCK2* and *ATF4* promoter regions (Figure 6L). These ARE-driven Luciferase reporter assays revealed that transcriptional expression of *CAT-ARE(#3)-Luc, G6PD-ARE(#9)-Luc* and *ATF4-ARE(#23)-Luc* was significantly activated by both Nrf1 and Nrf2 (Figure 6, M-Q). By contrast, Nrf2 alone also enabled transactivation of *G6PD-ARE(#10)-Luc, PCK2-ARE(#16)-Luc* and *ATF4-ARE(#19, #22* and *#24)-Luc* (Figure 6, M-Q), while only expression of *CAT-ARE(#5)-Luc* was up-regulated by Nrf1 (Figure 6M). However, expression of all three *PHGDH-ARE-Luc* reporters appeared to be unaffected by either Nrf1 or Nrf2 (Figure 6O).

## Discussion

In the previous study, we reported that knockout of *Nrf1α* leads to malignant proliferation and tumor metastasis [25, 30]. Such malignant growth and proliferation of cancer cells are also dictated by nutrient availability [38], because they require large amounts of nutrients intaked. On this basis, we herein discover that glucose starvation can prevent malignant proliferation of wild-type (*Nrf1/2*^*+/+*^) hepatocellular cancer cells and also causes cell death of *Nrf1α*-deficient hepatoma. This is fully consistent with the therapeutic strategy against cancer by its nutrients limiting [39].

It is, to our surprise, that glucose starvation leads to rapid cellular death of *Nrf1α*^−*/*−^ cells within 12 h, albeit with abberrant accumulation of Nrf2 in this difficient cells, whereas *Nrf2*^−*/*−^ cells manifest a strong resistance to the lethiality of glucose deprivation, even though Nrf1 is down-regulated. This finding demonstrates that both Nrf1 and Nrf2 may be disparately involved in setting the thresholds of distinct cellular patho-physiological (e.g., redox metabolic) responses. This notion is also supported by the evidence showing a small number of wild-type *Nrf1/2*^*+/+*^ cell deaths after glucose starvation for 24 h. Further examinations of cell death induced by glucose deprivation reveal that it is different from classical caspase-activated apoptosis, necroptosis, ferroptosis and autophagy. Notably, similar phenomenon was also observed after glucose starvation of other cell lines [40]. Contrarily, it is inferable that abnormal survival of *Nrf1α*^−*/*−^ cells are, in its malignant growth and proliferation state, maintained by a highly energy-consuming mechanism so as to require its ever-incrementing amounts of glucose and other nutrients. By contrast, the energy-consumption of *Nrf2*^−*/*−^ cells could be reduced to a considerable lower level than that of *Nrf1/2*^*+/+*^ cells. Thus, the putative demand for glucose is positively correlated with the subcutaneous tumorigenicility of distinct cancer xenografts in nude mice, as reported by Qiu et al [30]. Indeed, this is also corroborated by the previous finding that glucose uptake is substantially increased by silencing of *Nrf1* in pancreatic isle β-cells [29].

Further insights into acute death of glucose-starved *Nrf1α*^−*/*−^ cells have unveiled that such a strong lethality should be ascribed to severe endogenous oxidative stress and damage accumulation. This notion is substantiated by several lines of experimental evidence as followed. Firstly, accumulation of ROS in *Nrf1α*^−*/*−^ cells, albeit with hyper-active Nrf2, is significantly augmented by glycose deprivation, as accompanied by depletion of GSH. These, together, result in a striking increment of the GSSG/GSH ratio during glycose starvation of *Nrf1α*^−*/*−^ cells, in order to disrupt the intracellular redox balance and/or relevant signaling controls, as described by other groups [41, 42]. Secondly, death of *Nrf1α*^−*/*−^ cells induced by glucose starvation can be effectively prevented by both NAC and catalase, but not DHA. Similar results were obtained form treatment of other cell lines with NAC and catalase to rescue its glucose starvation-induced death [43]. Thirdly, it is found that the GSSG/GSH ratio of *Nrf2*^−*/*−^ cells is, conversely, diminished and even abolished during glycose starvation. Furtherly, silencing of Nrf2 in *Nrf1α*^−*/*−^ cells can also enable them to be alleviated from the cytotoxic ROS accumulation, so that glucose starvation-induced death of *Nrf1α*^−*/*−^ cells is sufficiently rescued by Nrf2 knockdown. Such surprising result implies that Nrf2 may also contribute to ROS production in *Nrf1α*^−*/*−^ cells under basal and glucose deprivation conditions, albeit it has been accepted as a master regulator of antioxidant cytoprotective genes [44]. This is also supported by the finding that *Nrf1α*^−*/*−^ cells are maintained at higher ROS levels in almost unstressed conditions, while *Nrf2*^−*/*−^ cells are preserved at relatively lower ROS levels than those of wild-type *Nrf1/2*^+*/*+^ cells (in this study). Fourthly, glucose starvation-induced expression of *PSMB5, PSMB6* and *PSMB7* (encoding the core proteasomal β5, β1, and β2 subunits) is substantially suppressed or even abolished in *Nrf1α*^−*/*−^ cells, besides their down-regulated basal expression. Such proteasomal dysfunction is likely to contribute to abberrant accumulation of oxidative damaged proteins (inclusing Nrf2) in *Nrf1α*^−*/*−^ cells. Together, our evidence demonstrates that Nrf1 is more potent than Nrf2 at mediating the cytoprotective responses against cytototic effects of glucose deprivation, and other *bona fide* cellular stressors, such as tunicamycin alone or plus *tert*-butylhydroquinone.

In-depth insights into the endogenous molecular basis for oxidative stress, contributing to the lethality of glucose deprivation, unravel that dysfunctional redox defense systems, along with altered redox signaling, are deteriorated in glucose-starved *Nrf1α*^−*/*−^ cells. In fact, we found that, though a large amount of ROS accumulation lead to acute death in *Nrf1α*−*/*− cells, but conversely, a relatively low concentration of ROS is also accumulated, facilitating maintenance of the malignant *Nrf1α*^−*/*−^ cell growth and proliferation. This finding is in full agreement with the double-edged effects of ROS, acting as two distinct and even opposing players in cell growth, differentiation, progression and death [45]. It is known that distinct high concentrations of ROS are involved in a variety of pathological processes, including cancer, ischemia, and immune and endocrine system deficiencies [45, 46]. The excessive ROS can also induc cell death by promoting the intrinsic apoptotic pathway [47]. Nonetheless, the low concentration of ROS are indispensable for various physiological processes, such as signal transduction and immune responses [46]. Such physiological ROS levels should be maintained in a steady-state by the homeostatic redox controls, including antioxidant defense systems, which comprise SOD, GPX1, GSR, CAT and other redox proteins. As a result, the intracellular superoxide anions, arising from aggressive mitochondrial metabolism [48] and other sources (as illustrated in Figure 3A), are converted by SOD to H_2_O_2_ and oxygen, and then H_2_O_2_ is decomposed by CAT into water and oxygen [49, 50]. Furtherly, oxidized glutathione (GSSG) can be reduced by GSR to yield the sulfhydryl GSH [51]. GPX1 catalyzes the reduction of H_2_O_2_ and organic hydroperoxides by glutathione, so as to protect cells against oxidative damage [52]. In this study, we demonstrate that basal expression of *CAT* and *GPX1* is evidently up-regulated in *Nrf1α*^−*/*−^ cells, but substantially suppressed by glucose starvation. By contrast, basal expression of *GSR* and *SOD1* is obviously down-regulated in *Nrf1α*^−*/*−^ cells, and further diminished by glucose deprivation. In addition, differential decreases of *TRX1, TRX2* and *SOD2* occur only after glucose deprivation. Overall, dysfunctions of redox signaling controls and/or antioxidant defense systems are aggravated by glucose starvation of *Nrf1α*^−*/*−^ cells, which contributes to the cellular lethality of this stress, albeit accumulation of hyper-active Nrf2. However, an exception to this is that NADPH oxidase 4 (NOX4), as a ROS-producing source, may also be co-regulated by Nrf1 and Nrf2, based on the evidence that basal and starvation-stimulated expression of NOX4 are significantly down-regulated in either *Nrf1α*^−*/*−^ or *Nrf2*^−*/*−^ cells.

To ameliorate the severe endogenous oxidative stress induced by glucose deprivation and thus facilitate survival and proliferation of cancer cells, they tend to redistribute those intermediates from glycolysis and gluconeogenesis to other metabolic pathways (e.g., PPP and SSP, in Figures 4A & 5A), particularly under glucose starvation. As stated by [53-55], the aerobic glycolysis, as a distinctive metabolic pattern of cancer cells from normal cells, provides a lot of intermediates for pentose phosphate pathway (PPP), gluconeogenesis and serine-to-glutathion synthesis pathway. Thus, this can enable cancer cells to reduce products of ROS from mitochondria and other subcellular compartments, but also enhance generation of NADPH and GSH from PPP and glutathion synthesis, respectively, so that both ratios of NADPH/NADP^+^ and GSH/GSSG are restored to a newly redox-balanced level. However, we here found that such altered glucose metabolism pathways are dysregulated in *Nrf1α*^−*/*−^ cells, and also deteriorated by glucose starvation, leading to the starved cell death. Among them mainly include increased glucose uptake, modestly reduced glycolysis, dysfunction of gluconeogenesis, PPP and SSP. As a matter of fact, glucose deprivation results in an abject failure of glucose uptake and ensuing glycolysis. Thereby, this confers gluconeogenesis to be gainig crucial importance, because gluconeogenesis is a potent alternative source of biosynthetic precursors under glucose deprivation, albeit its intermediates are shared from glycolytic pathways. Herein, we have proposed a conceptual mode (as illustrated in Figure 6, R-T), based on the evidence that glucose starvation of *Nrf1α*^−*/*−^ cells caused significant decreases in both abudances of PCK2 and PCK1 (as key rate-limiting enzymes of gluconeogenesis), leading to an enhancement of the starved cell death, but similar results were not obtained from glucose-starved *Nrf1/2*^*+/+*^ and *Nrf2*^−*/*−^ cells. Conversely, *de facto* transcripitional expression of *PCK2* in *Nrf1/2*^*+/+*^ or *Nrf2*^−*/*−^ cells was strikingly activated by glucose deprivation. Similar up-regulation of *PCK2* by low glucoses was also determined in A549 and H23 lung cancer cells, but its interferance and inhibition also significantly enhanced their apoptosis induced by glucose deprivation [56]. This notion is further corroborated by another evidence showing that PCK2, but not PCK1, is highly expressed in different cancer cell lines [54, 57].

The resulting intermediates of gluconeogenesis in glucose-starved cancer cells are allowed for divertion to enter the PPP and serine-to-glutathione synthesis pathways, so as to restore the intracellular redox (e.g., NADPH/NADP^+^ and GSH/GSSG) balances. Herein, we found that basal expression of *G6PD*, but not *PGD* (as two key enezymes to catalyze generation of NADPH), was up-regulated in *Nrf1α*^−*/*−^ cells, but glucose deprivation caused significant decreases of both expression in the exacerbated cellular death process. Similar decreased PPP flux, as accompaned by reduced NADPH levels and instead increased oxidative stress, was approved as a major fatal cause of the lethality of glucose starvation, because cell death was also accelerated by inhibiting G6PD [32]. However, it is full of curiosity that basal and glucose deprivation-stimulated expression levels of *G6PD* and *PGD* were substantially down-regulated by *Nrf2*^−*/*−^ cells to be considerable lower than those of *Nrf1α*^−*/*−^ cells, but rather *Nrf2*^−*/*−^ cells displayed a strong resistance to the lethality of glucose starvation. Such paradoxical observations indicate that other mechanisms, beyond PPP, are also involved in the response to glucose deprivation and its lethal cellular process. Thereby, distinct intracellular energy demands of ATP to determine cell survival or death decisions are also investigated, revealling that its basal ATP products are substantially augmented in *Nrf1α*^−*/*−^ cells, but abruptly repressed by glucose deprivation for 12 h to a much lower level than that of *Nrf1/2*^*+/+*^ cells. By contrast, basal ATP levels of *Nrf2*^−*/*−^ cells are markedly diminshed to the lowest level, and also almost unaffected by glucose deprivation. Such disparate energy-consuming demands of between *Nrf1α*^−*/*−^ and *Nrf2*^−*/*−^ cell lines dictate their survival or death decisions to be made, depending on gluconeogenesis and other nutrient sources in particular glucose deprivation conditions. In addition, we also discover that abundances of AMPK (as a key regulator of energy metabolism [34]) and its phosphorylated AMPK^Thr172^ (leading to its activation responsible for maintaining redox metabolic homeostasis [58]) are significantly suppressed by glucose starvation of *Nrf1α*^−*/*−^ and *Nrf2*^−*/*−^ cells, but with distinct (decreased or increased) ratios of phospho-AMPK^Thr172^/AMPK. Altogether, inactivation of AMPK to reduce ATP products is inferable as a main cause of blocking energy supply for glucose-starved *Nrf1α*^−*/*−^ cells, which results in a large number of these cell deaths occurring after 12 h of glucose deprivation.

Notably, serine biosynthesis is a vital turning point for glucose metabolism; its one hand provides an intermediate for anabolism, while its another hand directly affect cellular antioxidant capacity to generate cysteine and glutathione (as illustrated in Figure 5A, [59, 60]). However, the effects of glucose limitation on serine-to-glutathione synthesis in cancer cells are, for the first time, determined here. From all this, we discovered that all seven key genes (i.e., *PHGDH, PSAT1, PSPH, CBS, CTH, GCLC* and *GCLM*) for rate-limiting *de novo* serine-to-glutathione biosynthesis, along with *ATF4* (as a putative upstream regulator of serine synthesis [35]) and two antixidant genes (*HO-1* and *NQO1*), are substantially induced by glucose withdrawal from *Nrf1/2*^*+/+*^ cells. Among them, basal expression levels of *PHGDH, PSAT1, PSPH* and *CBS* were evidently up-regulated in *Nrf1α*^−*/*−^ cells, but unaffected or partially reduced by glucose deprivation. However, no significant changes in basal expression of these genes were observed in *Nrf2*^−*/*−^ cells, but they were still induced by glucose starvation, as compared to those of *Nrf1/2*^*+/+*^ cells. Such fatal defects of *Nrf1α*^−*/*−^, but not *Nrf2*^−*/*−^, cells in the serine synthesis and ensuing transsulfuration to yield cysteine demonstrate that Nrf1α plays a dominant regulator in the successive processes. By contrast, Nrf2 is a master regulator of GSH biosynthesis (*GCLC* and *GCLM*) and antixidant responsive genes (*HO-1* and *NQO1*) to glucose deprivation, albeit Nrf1 is also involved in controlling *GCLC* expression.

In summary, glucose deprivation induces conversion of Nrf1 glycoprotein and then proteolytic processing to give rise to its mature cleaved CNC-bZIP factor, in order to transcriptionally regulate distinct target gene expression. Besides, Nrf1-target proteasomal expression is also required for post-translational processing of key proteins (e.g., Nrf2), but its core proteasomal subunits are inhibited by glucose deprivation. Under this conditions, gluconeogenesis becomes a major alternative source of biosynthetic precursors, and its intermediates are shared from glycolytic pathway and then diverted into other synthetic pathways (e.g., PPP and SSP). Thereby, altered glucose metabolism and energy demands of *Nrf1α*^−*/*−^ cells are aggravated by glucose deprivation, even though hyper-active Nrf2 is accumulated by loss of *Nrf1α*. Of note, induction of serine-to-glutathione synthesis by glucose starvation is fatally abolished in *Nrf1α*^−*/*−^ cells, leading to severe endogenous oxidative stress and damage, as accompanied by up-regulation of antioxidant responsive genes by Nrf2. Such fatal defects of *Nrf1α*^−*/*−^, but not *Nrf2*^−*/*−^ cells in the redox metabolism reprogramming caused by glucose deprivation, lead rapidly to a large number of the former cell deaths, because the intracellular ROS are elevated, along with reduced ATP production (Figure 6, R-T). Overall, these demonstrate that Nrf1 should act as a dominant player in the redox metabolic reprogramming, and can also fulfill its cytoprotective response against fatal cytotoxity of glucose deprivation. Thus, existence of Nrf1 enables *Nrf2*^−*/*−^ cells to be endowed with a strong resistence to glucose starvation. Conversely, it is reasoned that Nrf2 cannot fulfill its cytoprotective function against severe oxidative stress and damage, though it serves as a master regulator of antioxidant response genes (e.g., *HO-1, NQO1*). Moreover, there exist certain inter-regulatory crosstalks between Nrf1 and Nrf2 in the redox metabolism reprogramming by co-regulating expression of some key responsive genes (e.g., *CAT, G6PD, PCK2, GCLC, ATF4*) to glucose deprivation.

## Materials and methods

### Cell culture and reagents

The human hepatocellular carcinoma HepG2 cells (*Nrf1/2*^*+/+*^) were obtained originally from the American Type Culture Collection (ATCC, Manassas, VA, USA). The fidelity was conformed to be true by its authentication profiling and STR (short tandem repeat) typing map (by Shanghai Biowing Applied Biotechnology Co., Ltd). On this base, *Nrf1α*^−*/*−^ and *Nrf2*^−*/*−^ (or *Nrf2*^−*/*−*ΔTA*^) were established by Qiu et al [30]. Before experimentation, they were maintained in Dulbecco’s modified Eagle’s medium (DMEM) containing 25 mmol/L high glucose, 10% (v/v) fetal bovine serum (FBS), 100 Units/ml penicillin-streptomycin, and cultured in a 37°C incubator with 5% CO_2_. In addition, it is noted that all key reagents and resources used in this study were listed in Table S1.

### Assays of cell death from glucose deprivation

Equal numbers (3×10^5^) of *Nrf1/2*^*+/+*^, *Nrf1α*^−*/*−^ and *Nrf2*^−*/*−^ cells were seeded in 6-well plates and allowed for growth in DMEM containing 25 mmol/L glucose and 10% FBS for 24 h. After reaching 80% of their confluence, they were then transfered to be cultured in fresh glucose-free DMEM for 6-24 h. Subsequently, the cell morphological changes in survival or death induced by glucose deprivation were observed by microscopy. These cells were then counted, after staining by trypan blue. In addition, *Nrf1α*^−*/*−^*+siNrf2* cells were also prepared here, and then subjected to the cell death assay, after they were allowed for glucose deprivation.

To rescue the cell death, fructose and mannose (25 mmol/L) were added to the glucose-free media, respectively. Addition of 2DG (10 mmol/L) was to restore the PPP in the absence of glucose. In order to identify distinct types of cell death, q-VD-OPH (10 μmol/L, as a pan-caspase inhibitor), Necrostatin-1 (100 μmol/L), Ferrostatin-1 (2 μmol/L) and 3-methyladenine (2 mmol/L, an autophagy inhibitor) were added to the glucose-free media, respectively. Futhermore, NAC (5-10 mmol/L), CAT (50 Units/ml) and DHA (100 μmol/L) were added in the glucose-free media to examine effects of antioxidants or pro-oxidants on cell death. After being incubated for 12-24 h, these cell morphological changes were visualized by microscopy, and the trypan-stained cells were also counted.

### Analysis of cell apoptosis and ROS by flow cytometry

Equal numbers (3 × 10^5^) of experimental cells (*Nrf1/2*^*+/+*^, *Nrf1α*^−*/*−^, *Nrf2*^−*/*−^, and *Nrf1α*^−*/*−^*+siNrf2*) were allowed for 24 h of growth in DMEM containing 25 mmol/L glucose and 10% FBS for. After reaching 80% of their confluence, these cells were subjected to glucose deprivation by being cultured in a fresh glucose-free medium for 12 h. Subsequently, these cells were incubated with Annexin V-FITC and propidium iodide (PI) for 15 min, before the cell apoptosis was analyzed by flow cytometry. Furtherly, the intracellular ROS levels was also determined, according to the instruction of ROS assay kit (Beyotime, Shanghai, China). The resulting data were further analyzed by the FlowJo 7.6.1 software.

### Assays of ATP levels and GSSG/GSH ratios

ATP levels are determined according to the instruction of enhanced ATP assay kit (Beyotime, Shanghai, China); GSSG/GSH ratios are determined according to the instruction of GSH and GSSG Assay Kit (Beyotime, Shanghai, China).

### Knockdown of *Nrf2* in *Nrf1α*^−*/*−^ cells by its siRNAs

A pair of double-stranded small RNAs targeting for the interference with *Nrf2* (i.e., siNrf2) were synthesized by TranSheep Bio Co.Ltd. (Shanghai, China). The oligonucleotide sequences are as follows: FW, 5′-GUAAGAAGCCAGAUGU UAAdTdT-3′; REV, 5′-UUAACAUCUGGCUUCUUACdTdT-3′. Subsequently, *Nrf1α*^−*/*−^ cells were transfected with 80 nmol/L of the *siNrf2* oligonucleotides in the mixture of Lipofectamine 3000 (Invitrogen, California, USA). Thereafter, the *siNrf2*-interferred cells were identified by RT-qPCR and Western blotting, before being experimentated.

### Real-time quantitative PCR analysis of gene expression

After all experimental cells reached 80% of their confluence, they were subjected to glucose starvation by being transferred in fresh glucose-free media. Their total RNAs were extracted after glucose deprivation for 0-12 h, before being subjected to the reactions with a reverse transcriptase to synthesize the first strand of cDNAs. Subsequently, the mRNA levels of examined genes in different cell lines were determined by RT-qPCR with the indicated pairs of their forward and reverse primers (as listed in TableS1). All the RT-qPCRs were carried out in the GoTaq real-time PCR detection systems by a CFX96 instrument (Bio-rad, Hercules, CA, USA). The resulting data were analyzed by the Bio-Rad CFX Manager 3.1 software (Bio-rad).

### Western blotting analysis of key functional proteins

After all experimental cells reached 80% of their confluence, they were subjected to glucose starvation by being transferred in fresh glucose-free media. After 6-12 h of glucose deprivation, their total proteins were extracted by lysis buffer (0.5% SDS, 0.04 mol/L DTT, pH 7.5) containing protease inhibitor cOmplete Tablets EASYpack or phosphatase inhibitor PhosSTOP EASYpack (each 1 tablet per 15 mL, Roche, Basel, Switzerland), and diluted in 6×SDS-PAGE sample loading buffer (Beyotime, Shanghai, China). Subsequently, total lysates were denatured immediately at 100°C for 10 min, before equal amounts of proteins were separated by SDS-PAGE gels containing 8-12% polyacrylamide and visualized by Western blotting with distinct primary antibodies (as listed in Table S1). Among included those antibodies against Nrf1; Nrf2, ATF4, GLUT1, GPX1, TRX1, TRX2, PRX1, CBS, HO1, GCLM, GCLC, GSR, SOD1, PCK1, PCK2 and G6PD, CAT, AMPK and p-AMPK^Thr172^, PHGDH, PSAT1, HK1, HK2, PSMB5, PSMB6 and PSMB7. In addition, β-Actin served as an internal control to verify amounts of proteins that were loaded in each of wells.

### Assays of ARE-driven luciferase reporter gene activity

The core ARE consensus sites within -4K-bp sequences to the transcription start sites (TSS) or extended to the translation initiation sites (TIS) of *ATF4, CAT, G6PD, PHGDH* and *PCK2* promoter regions were searched. Each of the core ARE and adjoining sequences was then inserted into the indicated site of pGL3-promoter vector. The fidelity of all resultant constructs was confirmed by sequencing with indicated primer pairs (Table S1). Subsequently, equal numbers (1.5 × 10^5^) of HepG2 cells were seeded in 12-well plates and allowed for 24-h growth in DMEM containing 25 mmol/L glucose and 10% FBS. After reaching 80% confluence, the cells were transfected with expression construct for human Nrf1 or Nrf2, along with each of the above *ARE-Luc* reporters plus pRL-TK (serves as an internal control). Approximately 24 hours after transfection, ARE-driven luciferase activity was measured by using the dual-luciferase reporter assay. The resulting data were calculated as fold changes (mean ± S.D, n=9), relative to the basal activity (at a given value of 1.0) obtained from transfection of cells with an empty pcDNA3.1 and each of ARE-driven reporter genes.

### Statistical Analysis

Statistical significances of fold changes in gene expression levels measured by RT-qPCR and luciferase assays were assessed by using the Student’s *t*-test. The data are shown as a fold change (mean ± S.D, n=9) and each of which represents at least three independent experiments, that were each performed in triplicate.

## Author contributions

Y.-P.Z. performed the most experiments with help of Z.Z., and collected relative data, except that Y.X. did the experimental analysis of the half-lives of Nrf1 and Nrf2. Y.-P.Z. also made draft of this manuscript with most figures and supplemental tables. Y.Z. designed and supervised this study, analyzed all the data, helped to prepare all figures with cartoons, wrote and revised the paper.

## Acknowledgments

We are greatly thankful to Drs. Lu Qiu (at Zhengzhou University, China) and Yonggang Ren (North Sichuan Medical College, Sichuan, China) for having estabolished those relevant cell lines used in this study. We also thank to Mr. Shaofan Hu and other members (at Chongqing University, China) for giving invaluable help with this work. Of note, the study was supported by the National Natural Science Foundation of China (NSFC, with a key program 91429305 and another project 81872336) awarded to Yiguo Zhang (at Chongqing University, China). This work is also, in part, funded by Sichuan Department of Science and Technology grant (2019YJ0482) to Dr. Yuancai Xiang (at Southwest Medical University, Sichuan, China).

## Figure legends

**Figure S1. Distinct cellular responses to glucose deprivation.**

(A,B) *Nrf1/2*^*+/+*^, *Nrf1α*^−*/*−^ and *Nrf2*^−*/*−^ cells had been starved, or not starved, in glucose-free media for 6 h, before their morphological observation by microscopy (*A*, with an original magnification of 200×) and counting of dead cells by trypan blue staining (*B*).

(C,D) The above experimental cells were or were not subjected to glucose starvation for 24 h, before morphological observation (*C*) and dead cell counting (*D*).

(E) Apoptosis of *Nrf1α*^−*/*−^ *+ siNrf2* cells, that had been glucose-starved for 12 h, were determined by Flow cytometry.

**Figure S2. Time-dependent effects of glucose deprivation on Nrf1 and Nrf2 with distinct half-lives.**

(A) HepG2 cells were, or were not, co-treated for 0-4 h in the glucose-free media to which cycloheximide (CHX, 50 µg/ml) alone or plus bortezomib (BTZ, 1 µmol/L resoved in DMSO) was added, before being harvested. The total lysates were resolved by SDS-PAGE gels containing 8% and 12% polyacrylamide, followed by Western blotting with antibodies against Nrf1 or Nrf2.

(B) The relative intensity of major protein bands representing Nrf1 and Nrf2 was quantified and shown graphically. The [X]_t_/[A]_0_ represent the relative amount of indicated protein at the indicated times ([A]_t_) was normalized to [A]_0_. The half-lives of the major Nrf1 and Nrf2 isoforms were calculated and shown herein.

**Table S1. The key reagents and resources used in this work.**

